# Nicotianamine synthase 2 is required for symbiotic nitrogen fixation in *Medicago truncatula* nodules

**DOI:** 10.1101/717983

**Authors:** Viviana Escudero, Isidro Abreu, Eric del Sastre, Manuel Tejada-Jiménez, Camile Larue, Lorena Novoa-Aponte, Jiangqi Wen, Kirankumar S. Mysore, Javier Abadía, José M. Argüello, Hiram Castillo-Michel, Ana Álvarez-Fernández, Juan Imperial, Manuel González-Guerrero

## Abstract

Symbiotic nitrogen fixation carried out by the interaction between legumes and diazotrophic bacteria known as rhizobia requires of relatively large levels of transition metals. These elements act as cofactors of many key enzymes involved in this process. Metallic micronutrients are obtained from soil by the roots and directed to sink organs by the vasculature, in a process participated by a number of metal transporters and small organic molecules that mediate metal delivery in the plant fluids. Among the later, nicotianamine is one of the most important. Synthesized by nicotianamine synthases (NAS), this non-proteinogenic amino acid forms metal complexes participating in intracellular metal homeostasis and long-distance metal trafficking. Here we characterized the *NAS2* gene from model legume *Medicago truncatula*. MtNAS2 is located in the root vasculature and in all nodule tissues in the infection and fixation zones. Symbiotic nitrogen fixation requires of *MtNAS2* function, as indicated by the loss of nitrogenase activity in the insertional mutant *nas2-1*, a phenotype reverted by reintroduction of a wild-type copy of *MtNAS2*. This would be the result of the altered iron distribution in *nas2-1* nodules, as indicated by X-ray fluorescence studies. Moreover, iron speciation is also affected in these nodules. These data suggest a role of nicotianamine in iron delivery for symbiotic nitrogen fixation.

**Significance Statement:** Nicotianamine synthesis mediated by MtNAS2 is important for iron allocation for symbiotic nitrogen fixation by rhizobia in *Medicago truncatula* root nodules.

## INTRODUCTION

Nitrogen is one of the main limiting nutrients in the biosphere, in spite of N_2_ abundance (Smil, 1999, Hoffman *et al.*, 2014). Nitrogenase is the only enzyme that can convert, fix, N_2_ into NH_3_ under physiological conditions, in an energy consuming process (Burk, 1934, Burgess and Lowe, 1996). This enzyme is only expressed by the small group of diazotrophic archaea and bacteria, some of them participating in symbiosis with other organisms (Boyd and Peters, 2013). Arguably, one of the best characterized symbiosis with diazotrophic bacteria is the one established between rhizobia and legumes (Brewin, 1991, Downie, 2014). This symbiosis is the basis for legume use in crop rotation strategies and their potential as an alternative to polluting and expensive synthetic nitrogen fertilizers (Johnson and Mohler, 2009, Mus *et al.*, 2016).

Symbiotic nitrogen fixation by the legume-rhizobia system is carried out in root nodules (Downie, 2014). These are differentiated organs that develop after a complex exchange of chemical signals between the symbionts (Oldroyd, 2013). Detection of the nodulation factors released by the rhizobia, triggers cell proliferation in the pericyle-inner cortex of the root to originate nodule primordia (Xiao *et al.*, 2014). As nodules grow, rhizobia from the root surface are directed by infection threads to the nodule cells (Gage, 2002). There, they are released in an endocytic-like process, originating pseudo-organelles known as symbiosomes (Roth and Stacey, 1989, Catalano *et al.*, 2006). Within the symbiosomes, rhizobia differentiate into bacteroids and express the enzymatic machinery required for nitrogen fixation (Kondorosi *et al.*, 2013). Nodule development follows either an indeterminate or a determinate growth pattern, based on whether they maintain an apical meristem to sustain growth (Vasse *et al.*, 1990). As this meristem allows for sustained growth in indeterminate nodules, four developmental zones appear: the meristematic region or Zone I; the infection-differentiation zone or Zone II, where rhizobia are released in the cell and start differentiating; the fixation zone or Zone III, where nitrogenase is active; and the senescent zone or Zone IV, where symbiosomes are degraded and nutrients recycled (Burton *et al.*, 1998). In addition, some authors define a transition interzone between Zones II and III (Roux *et al.*, 2014).

Nutrient exchange between the symbionts enables nitrogen fixation (Udvardi and Poole, 2013). Availability of fixed nitrogen forms in soils inhibits nodulation (Streeter, 1987). Similarly, low levels of photosynthates, phosphate or sulphate transfer from the host plant decrease nodulation and nitrogen fixation rates (Singleton and van Kessel, 1987, Valentine *et al.*, 2017, Schneider *et al.*, 2019). Transition metals such as iron, copper, zinc, or molybdenum are also critical for nodulation and nitrogen fixation as cofactors in many of the involved enzymes (González-Guerrero *et al.*, 2014, González-Guerrero *et al.*, 2016). This includes not only nitrogenase (Rubio and Ludden, 2005), but also NADPH-oxidases that participate in nodule signalling (Montiel *et al.*, 2016), leghemoglobin that maintains nodule O_2_ homeostasis (Appleby, 1984), high-affinity cytochrome oxidases providing energy to the bacteroids (Preisig *et al.*, 1996), as well as many enzymes involved in free radical control (Dalton *et al.*, 1998, Santos *et al.*, 2000, Rubio *et al.*, 2007). Consequently, deficiencies in the uptake of these nutrients or alterations in the metal delivery pathways lead to defects in nodulation and/or nitrogen fixation (Tang *et al.*, 1991, O’Hara, 2001, Senovilla *et al.*, 2018, Gil-Díez *et al.*, 2019).

To reach the bacteroids, metals must first cross from soil into the roots using the general mechanisms common to all dicots (Kobayashi and Nishizawa, 2012, Curie and Mari, 2017). Metal uptake is facilitated by soil acidification, the release of phenolics/coumarins and flavins, and cation reduction when required (Jain *et al.*, 2014). Metals are then introduced into the root epidermis and symplastically or apoplastically reach the root endodermis, to cross into the vasculature, and delivered to sink organs. In model legume *Medicago truncatula* metals are released from the vessels into the apoplast of the infection-differentiation zone of nodules (Rodríguez-Haas *et al.*, 2013). These nutrients will be introduced in rhizobia-infected cells and targeted to symbiosomes for nitrogen fixation. In recent years, many of the membrane transporters participating in metal transfer from the plant to the bacteroids have been identified. For instance, iron transfer to nitrogen-fixing cells is facilitated by plasma membrane iron uptake protein MtNramp1 (Tejada-Jiménez *et al.*, 2015), and its transport across the symbiosome membrane by MtSEN1 and MtFPN2 (Hakoyama *et al.*, 2012, Escudero *et al.*, 2019). However, little is known on how metals are sorted intracellularly and on the speciation of these elements.

Unlike alkali or alkali-earth elements, transition metals are not “free”, hydrated, in physiological solutions. Instead, they are bound to a plethora of organic molecules that maintain them soluble under different pH, prevent metal-catalysed production of free radicals in Fenton-style reactions, and avoid mis-metallation of enzymes (Finney and Halloran, 2003, Rellán-Álvarez *et al.*, 2008, Flis *et al.*, 2016). Systematic studies of the nature of these chemical species in the sap of model plants have revealed the importance of citrate and nicotianamine in this role (von Wiren *et al.*, 1999, Durrett *et al.*, 2007, Roschzttardtz *et al.*, 2011, Schuler *et al.*, 2012). Citrate is the main iron chelator in xylem and facilitates iron delivery across symplastically disconnected tissues (Durrett *et al.*, 2007, Rellán-Álvarez *et al.*, 2010, Roschzttardtz *et al.*, 2011). It has also been associated with iron trafficking to nodules (LeVier *et al.*, 1996). Citrate efflux proteins LjMATE1 and MtMATE67 are required for iron allocation to nodules and contribute to nitrogen fixation (Takanashi *et al.*, 2013, Kryvoruchko *et al.*, 2018). Citrate efflux is also important for iron delivery to bacteroids, as indicated by the symbiosome localization of nodule-specific protein MtMATE67 (Kryvoruchko *et al.*, 2018).

Nicotianamine is also an important player in plant metal homeostasis. This molecule is a non-proteinogenic amino acid synthesized by nicotianamine synthases (NAS) from S-adenosyl methionine (Higuchi *et al.*, 1999). Nicotianamine-metal complexes mediate long-distance metal trafficking, particularly along the phloem, as well as participate in vacuolar metal storage (von Wiren *et al.*, 1999, Haydon *et al.*, 2012, Flis *et al.*, 2016). A nodule-specific NAS gene was identified in senescent nodules of *L. japonicus*, likely participating in the metal redistribution to the developing flowers and embryos, as orthologues do with older leaves (Hakoyama *et al.*, 2009, Schuler *et al.*, 2012). No such nodule-specific NAS gene can be found in transcriptomic databases from indeterminate type nodules, but tentative evidence shows that a *M. truncatula* NAS protein, MtNAS1, might be responsible for iron allocation to these organs (Avenhaus *et al.*, 2016). Here, we have characterized a second NAS protein, MtNAS2, identified in a screening of *M. truncatula Tnt1*-insertion mutants. This gene, although primarily expressed in roots, is important for metal allocation for symbiotic nitrogen fixation.

## RESULTS

### *Medicago truncatula Tnt1* line NF15101 phenotype is due to transposon insertion in *MtNAS2*

A search for metal-related symbiotic phenotypes of the mutants available at the Noble Research Institute *M. truncatula* Mutant Database showed NF15101 as one of the available mutants with a nitrogen fixation deficient phenotype. This line has 22 *Tnt1* insertions, 10 of which interrupted different *M. truncatula* genes (Supp. Table 1), *Medtr2g070310* among them. This gene encodes a protein with 52 % identity and 67 % similarity to *Arabidopsis thaliana* NAS2 protein, and consequently was renamed *MtNAS2*. *MtNAS2* was expressed at similar levels in roots from plants inoculated or non-inoculated with *Sinorhizobium meliloti* (Fig. 1A). Significantly, lower expression was detected in nodules, and no signal was detected in shoots from either inoculated or non-inoculated plants. *Tnt1* was inserted in position +760 of *MtNAS2*, interrupting the reading frame of its only exon and diminishing *MtNAS2* mRNA levels below our detection limit (Fig. 1B). As expected, NF15101, *nas2-1* in this report, had reduced biomass production in nitrogen fixation conditions (Fig. 2A, B). While nodule development and nodule number were not significantly altered in *nas2-1* compared to wild type (Fig. 2C, D, Supp. Fig. 1), nitrogenase activity was reduced three-fold in *nas2-1* plants (Fig. 2E). No significant differences in nicotianamine content were observed between wild-type and mutant plants (Supp. Fig. 2). The *nas2-1* phenotype was reverted when a wild-type copy of *MtNAS2* regulated by its own promoter was reintroduced in *nas2-1* (Fig. 2). The data indicate that among all the *Tnt1* insertions, loss of *MtNAS2* function was determinant for the reduction of nitrogenase activity and overall growth alterations.

**Figure 1.**
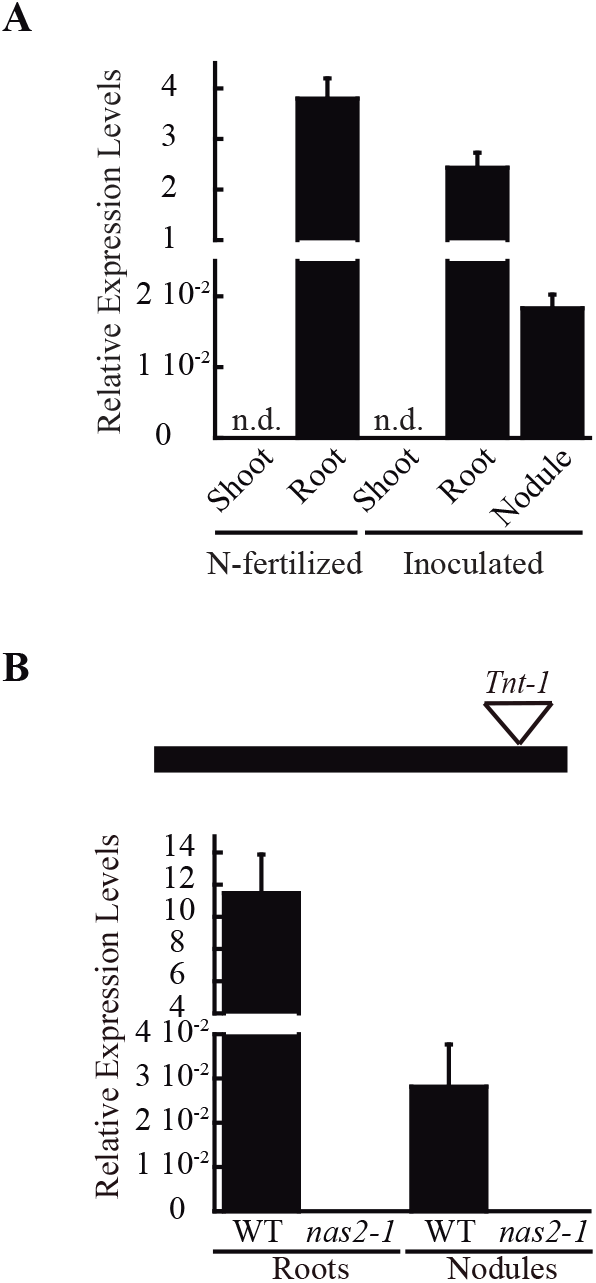
*MtNAS2* is expressed in roots and nodules of *M. truncatula*. (A) Expression data was normalized to the expression of ubiquitin conjugating enzyme E2 gene (*Medtr7g116940*) as standard. Data are the mean ± SE of three independent experiments with 4 pooled plants. (B) *Tnt1* insertion in the only exon of *MtNAS2* causes loss of *MtNAS2* transcripts. Expression was determined in 28 dpi roots and nodules of wild-type (WT) and *nas2-1* plants. Data was relativized to the expression of ubiquitin conjugating enzyme E2 (*Medtr7g116940*) and expressed as mean ± SE of three independent experiments with 4 pooled plants.

**Figure 2.**
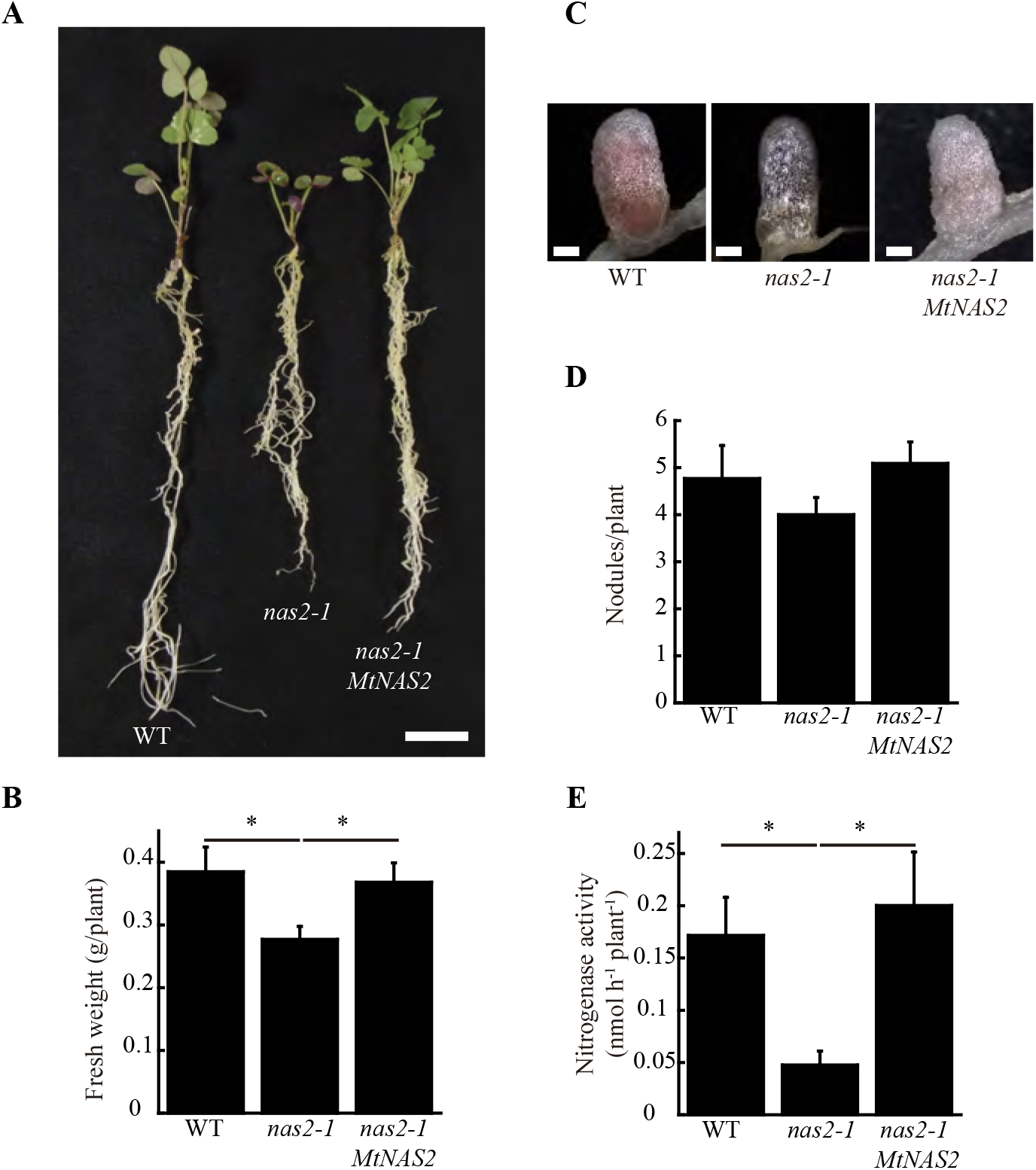
*MtNAS2* is required for nitrogen fixation. (A) Growth of representative wild-type (WT), *nas2-1*, and *nas2-1* plants transformed with *MtNAS2* controlled by its own promoter (*nas2-1 MtNAS2*). Bar = 1.5 cm. (B) Fresh weight of WT, *nas2-1*, and *nas2-1 MtNAS2* plants. Data are the mean ± SE of at least 9 transformed plants. (C) Detail of representative nodules of WT, *nas2-1*, and *nas2-1 MtNAS2* plants. Bars = 500 μm. (D) Number of nodules in 28 dpi WT, *nas2-1*, and *nas2-1 MtNAS2* plants. Data are the mean ± SE of at least 9 transformed plants. (E) Nitrogenase activity in 28 dpi nodules from WT, *nas2-1*, and *nas2-1 MtNAS2* plants. Acetylene reduction was measured in duplicate from three sets of three-four pooled plants. Data are the mean ± SE. * indicates statistically significant differences (p<0.05).

### *MtNAS2* is not required for plant growth under non-symbiotic conditions

To determine whether the symbiotic phenotype of *nas2-1* was the result of additional physiological processes being affected, these plants and their controls were grown in the same conditions as above, but supplemented with ammonium nitrate in the nutrient solution to compensate for the lack of rhizobial inoculation. In these conditions, no significant differences were found in plant growth, biomass production, or chlorophyll content between wild-type and *nas2-1* plants (Fig. 3). Considering the role of nicotianamine in plant iron homeostasis (von Wiren *et al.*, 1999, Inoue *et al.*, 2003) and the added pressure of symbiotic nitrogen fixation on iron nutrition (Terry *et al.*, 1991), *nas2-1* phenotype was also studied under non-symbiotic, low-iron conditions (no iron added to the nutrient solution). Low-iron supply did not lead to different growth between control and *nas2-1* plants (Supp. Fig. 3).

**Figure 3.**
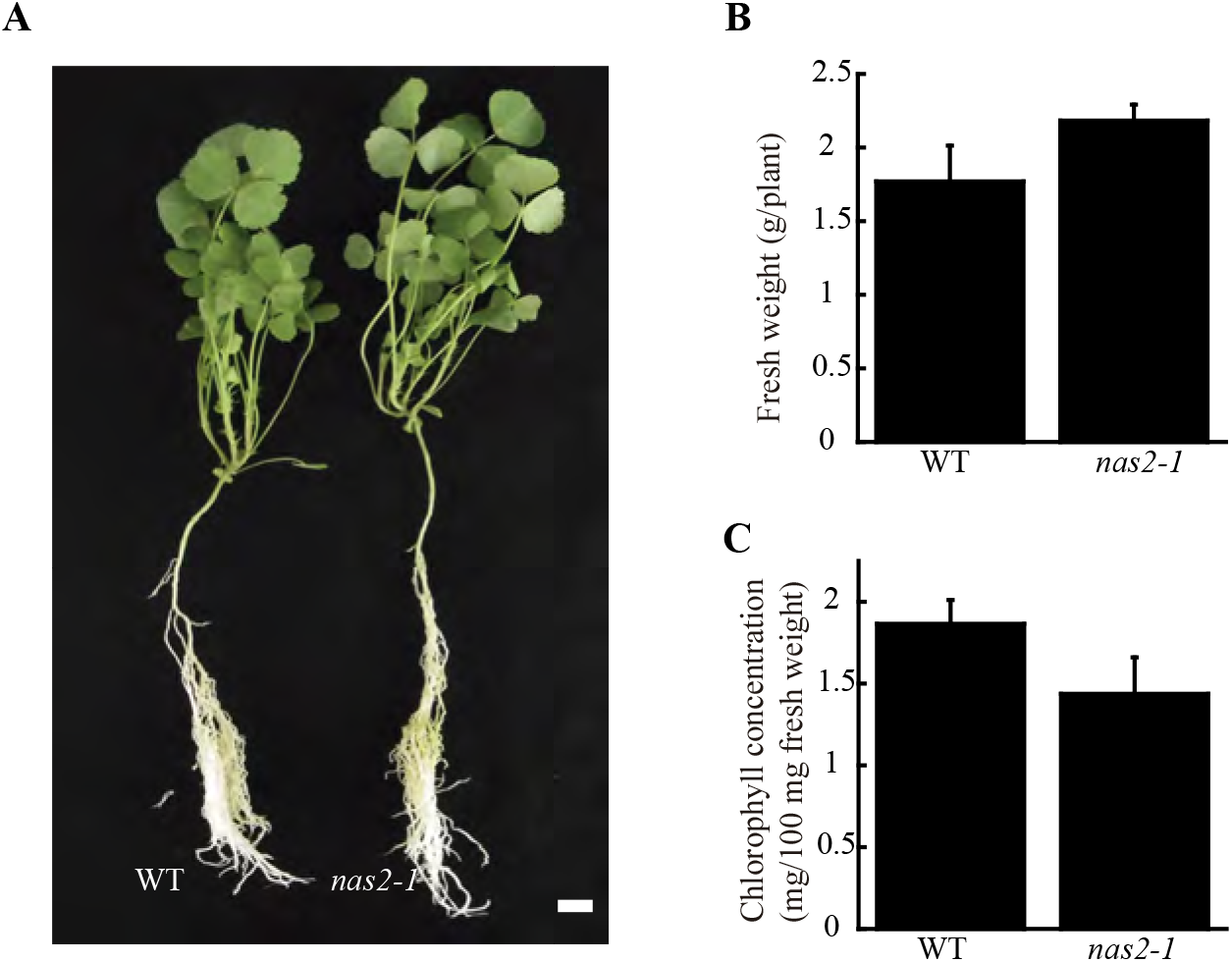
*MtNAS2* is not required for plant growth under non-symbiotic conditions. (A) Growth of representative wild-type (WT) and *nas2-1* plants when watered with a nutrient solution supplemented with ammonium nitrate and not inoculated with *S. meliloti*. Bar = 1.5 cm. (B) Fresh weight of WT and *nas2-1* plants. Data are the mean ± SE of at least 5 plants. (C) Chlorophyll concentration of wild-type and *nas2-1* plants. Data are the mean ± SE of two sets of 5 pooled plants.

### *MtNAS2* is expressed in the xylem parenchyma in roots and in the nodule fixation zone

The physiological role of *MtNAS2* is determined by its differential tissue and cellular expression. To establish the gene tissue expression, *M. truncatula* plants were transformed with a binary vector containing the *MtNAS2* promoter region driving the *β*-glucuronidase (*gus*) gene transcription and GUS activity visualized using X-Gluc. *MtNAS2* was expressed in roots and nodules (Fig. 4A), in agreement with the transcript data (Fig. 1). Longitudinal section of the nodules showed GUS activity in cells in the interzone and fixation zone of the nodule (Fig. 4B). Nodule cross-sections showed expression in all nodule tissues (Fig. 4C). In roots, *MtNAS2* promoter was active in vasculature cells (Fig. 4D).

**Figure 4.**
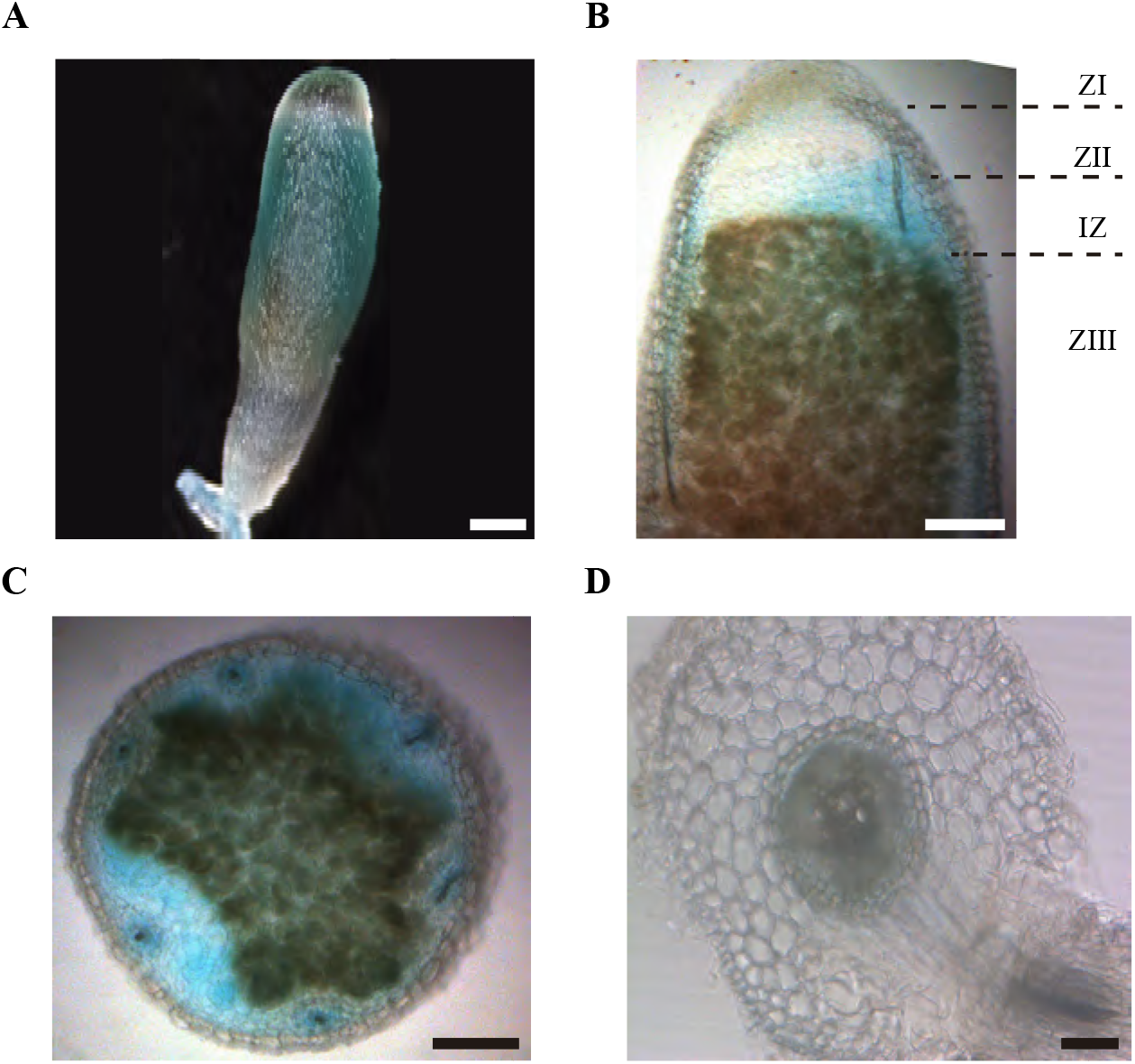
*MtNAS2* is expressed in the root vasculature and in the interzone, zone III, and vessels in nodules. (A) GUS staining of 28 dpi *M. truncatula* roots and nodules expressing the *gus* gene under the control of *MtNAS2* promoter region. Bar = 200 μm. (B) Longitudinal section of a GUS-stained 28 dpi *M. truncatula* nodule expressing the *gus* gene under the control of *MtNAS2* promoter region. ZI indicates Zone I; ZII, Zone II; IZ, Interzone; and ZIII, Zone III. Bar = 200 μm. (C) Cross section of a GUS-stained 28 dpi *M. truncatula* nodule expressing the *gus* gene under the control of *MtNAS2* promoter region. Bar = 200 μm. (D) Cross section of a GUS-stained 28 dpi *M. truncatula* root expressing the *gus* gene under the control of *MtNAS2* promoter region. Bar = 50 μm.

Supporting the gene expression results, immunolocalization of HA-tagged MtNAS2 under control of its own promoter showed that the protein was located in cells neighbouring the interzone and fixation zone (Fig. 5A). At higher magnification, we could observe that MtNAS2-HA had a homogenous distribution within the cells, and it did not seem to cluster in any particular location (Fig. 5B). Analysis of nodule vasculature showed MtNAS2-HA in endodermal cells (Fig. 5C). However, in the root vasculature, MtNAS2-HA was detected in small cells associated to the xylem, (Fig. 5D). Controls were carried out to ensure that the data did not stem from autofluorescence (Supp. Fig 4).

**Figure 5.**
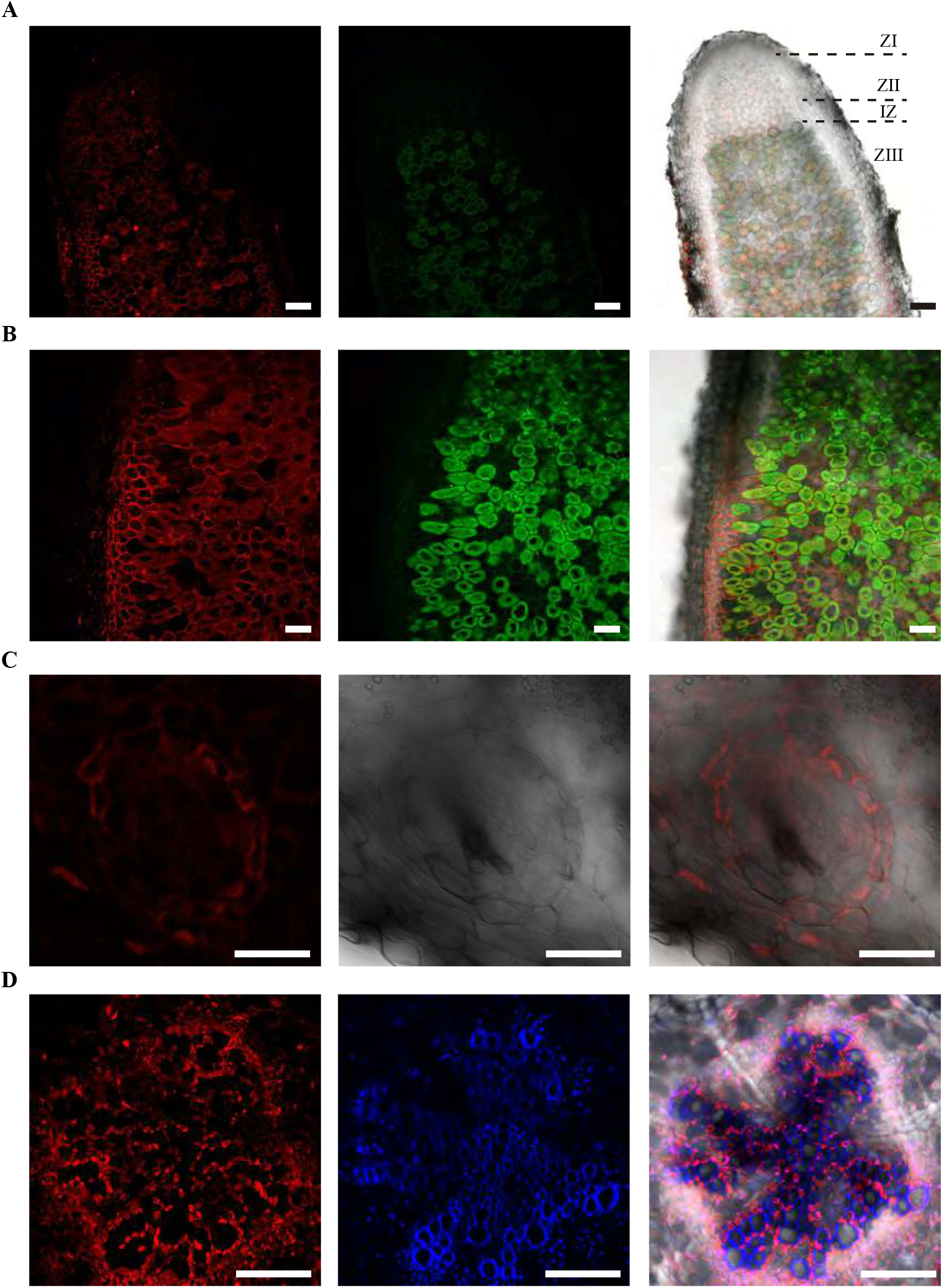
MtNAS2 is located in the nodule core cells, in the endodermis of the nodule vessels, and in cells surrounding the xylem in the root vasculature. (A) Longitudinal section of a 28 dpi *M. truncatula* nodule expressing *MtNAS2-HA* under its own promoter. The three C-terminal HA epitopes were detected using an Alexa594-conjugated antibody (red, left panel). Transformed plants were inoculated with a GFP-expressing *S. meliloti* (green, middle panel). Both images were overlaid with the transillumination image (right panel). ZI indicates Zone I; ZII, Zone II; IZ, Interzone; and ZIII, Zone III. Bars = 100 μm. (B) Detail of the zone III of a 28 dpi *M. truncatula* nodule expressing *MtNAS2-HA* under its own promoter. Left panel corresponds to the Alexa594 signal used to detect the HA-tag, middle panel corresponds to the GFP channel showing *S. meliloti*, and the two were overlaid with the bright field channel in the right panel. Bars = 50 μm (C) Cross section of a nodule vessel from a 28 dpi *M. truncatula* nodule expressing *MtNAS2-HA* under its own promoter. Left panel corresponds to the Alexa594 signal used to detect the HA-tag, middle panel corresponds to the bright field channel showing *S. meliloti*, and the two were overlaid in the right panel. Bars = 50 μm. (D) Cross section from a 28 dpi *M. truncatula* root expressing *MtNAS2-HA* under its own promoter. Left panel corresponds to the Alexa594 signal used to detect the HA-tag, middle panel corresponds to autofluorescence signal of xylem, and the two were overlaid with the bright field channel in the right panel. Bars = 100 μm.

### *MtNAS2* is required for efficient metal allocation for symbiotic nitrogen fixation

Nicotianamine is required for metal allocation from source to sink tissues (Schuler *et al.*, 2012). Alterations in nicotianamine synthesis typically lead to reduced metal delivery to sink tissues. To determine whether this was the case for *nas2-1*, iron, copper and zinc levels in roots, shoots, and nodules from 28 days-post-inoculation (dpi) plants were determined. No significant changes in these levels were observed (Fig. 6A). However, metal allocation might be altered while not affecting total nodule metal content. To assess this possibility, synchrotron-based X-ray fluorescence studies were carried out to determine iron distribution in *nas2-1* compared to wild type (Fig. 6B). These experiments showed that iron distribution was altered in *nas2-1* mutants. To further confirm that mutation of *MtNAS2* affected iron distribution in nodules as a consequence of changes of iron speciation, X-ray Absorption Near-Edge Spectroscopy (XANES) analyses of iron speciation in the different nodule developmental zones were carried out (Fig. 6C). Principal component analyses of these spectra showed that the iron complexes in the fixation zone were quite different (Fig. 6D). Fitting of the obtained spectra to known standards showed that the proportion of Fe-S complexes had a dramatic drop in *nas2-1* compared to wild-type plants, while the proportion of O/N complexes with iron had a larger increase (Table 1).

**Figure 6.**
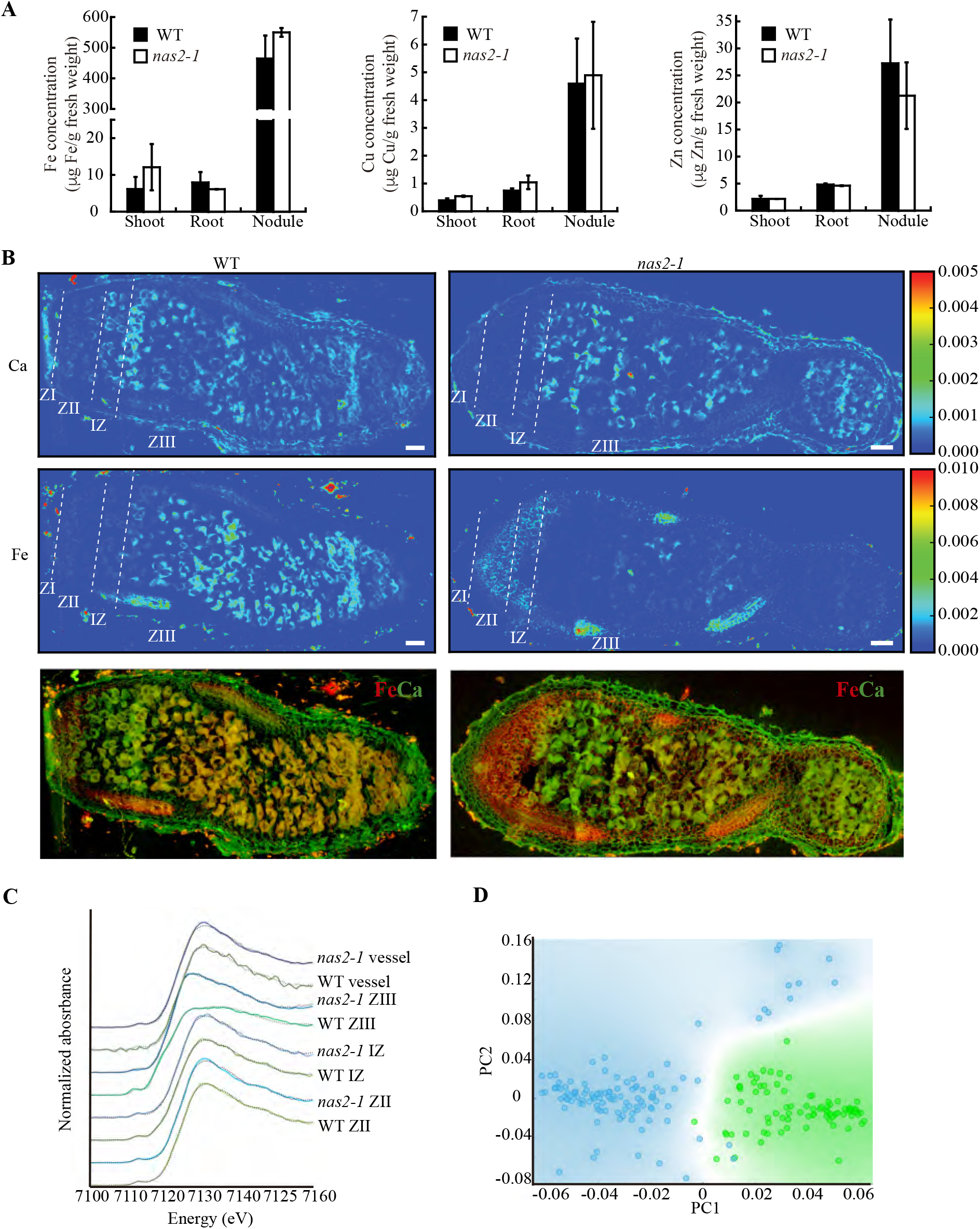
*MtNAS2* is required for iron distribution and speciation in nodules. (A) Iron (left panel), copper (middle panel), and zinc (right panel) concentration in shoots, roots, and nodules from 28 dpi wild-type (WT) and *nas2-1* plants. Data are the mean ± SE of three sets of three-four pooled organs. (B) Synchrotron-based X-ray fluorescence images of WT (left panels) or *nas2-1* (right panels) showing calcium (top panels) or iron (center panels) distribution in 28 dpi nodules. Lower panels are the overlaid iron and calcium distribution (iron is indicated in red and calcium in green). ZI indicates Zone I; ZII, Zone II; IZ, Interzone; and ZIII, Zone III. Bars = 100 μm. (C) XANES spectra obtained from different regions of WT and *nas2-1* nodules. (D) Decomposition of the Zone III signal into its two principal components.

**Table 1:** Iron speciation (%) in WT and *nas2-1* nodules. n.d. = not detected.

| Region | WT |  |  |  | *nas2-1* |  |  |  |
| --- | --- | --- | --- | --- | --- | --- | --- | --- |
|  | Fe(II)-<br>S | Fe(II)-<br>O/N | Fe(III)-<br>O | R<sup>2</sup> | Fe(II)-<br>S | Fe(II)-<br>O/N | Fe(III)-<br>O | R<sup>2</sup> |
| ZII | 4 | 12 | 83 | 00004 | n.d. | 14 | 86 | 0.0008 |
| IZ | 13 | 19 | 69 | 0.0004 | 5 | 17 | 78 | 0.0006 |
| ZIII | 50 | 24 | 27 | 0.001 | 14 | 49 | 37 | 0.0005 |
| Vessels | n.d. | 22 | 79 | 0.0008 | n.d. | 22 | 78 | 0.001 |

## DISCUSSION

Metallic micronutrient delivery to nodules is essential for symbiotic nitrogen fixation, as they are cofactors in many of the involved enzymes (Brear *et al.*, 2013, González-Guerrero *et al.*, 2014). In recent years, studies have shown how metals are exported to the apoplast in the infection/differentiation zone of *M. truncatula* nodules (Rodríguez-Haas *et al.*, 2013), and transmembrane transporters introduce metals into rhizobia-infected cells (Tejada-Jiménez *et al.*, 2015, Abreu *et al.*, 2017, Tejada-Jiménez *et al.*, 2017, Senovilla *et al.*, 2018), or deliver iron to the bacteroids (Escudero et al., 2019). In this transport, citrate participates in maintaining iron solubility in the apoplast, and as the preferred iron source for bacteroids (Moreau *et al.*, 1995, LeVier *et al.*, 1996, Kryvoruchko *et al.*, 2018). Here we show that nicotianamine synthesis is also important for correct iron allocation to *M. truncatula* nodules.

Interrupting *MtNAS2* expression with a transposon insertion led to reduced plant growth in symbiotic conditions, a consequence of lower nitrogenase activity. Although several genes were affected in the studied *Tnt1* line, reintroduction of a wild-type copy of *MtNAS2* was sufficient to restore wild-type growth. Consequently, the mutation of this gene was mainly responsible for the observed phenotype. While important under symbiotic conditions, *MtNAS2* seemed to be playing a secondary role when plants were not inoculated but watered with an ammonium nitrate-supplemented nutrient solution instead. This is in contrast to the substantially higher expression levels of *MtNAS2* in roots than in nodules. This observation would suggest a predominant role in nicotianamine synthesis in roots. However, studies in *A. thaliana* reveal the existence of a high redundancy rate in the NAS family, where a quadruple *nas* mutant was required to observe a substantial phenotype, including limited growth (Klatte *et al.*, 2009). Similarly, no significant changes in nicotianamine content were observed in single *nas A. thaliana* lines, as neither was observed in *M. truncatula nas2-1*. Two possible causes might explain the symbiosis-specific phenotype of *nas2-1* plants. One of them is that MtNAS2 would be required to compensate for the enhanced iron requirements of nodulated plants. This additional nutritional pressure would trigger the observed *nas2-1* phenotype. If so, we should have also observed a similar phenotype when plants were watered with an iron-restricted nutrient solution, which has been shown in the past to elicit the iron deficiency response in *M. truncatula* (Andaluz *et al.*, 2009, Tejada-Jiménez *et al.*, 2015). However, this was not observed. Alternatively, in a more parsimonious mechanism, neofunctionalization of pre-existing genes during the development of symbiotic nitrogen fixation might have led to the loss of functional redundancy. Similar observations have been made when studying other *M. truncatula* metal homeostasis genes that, although expressed in roots and in nodules, exhibit phenotypes limited to nodulation and nitrogen fixation (Tejada-Jiménez *et al.*, 2015, Abreu *et al.*, 2017, León-Mediavilla *et al.*, 2018).

In roots, *MtNAS2* was expressed at high levels in xylem parenchyma cells, similarly to rice *NAS2* (Inoue *et al.*, 2003). Vascular localization of NAS proteins is not unusual, since they have been associated to long distance metal trafficking (von Wiren *et al.*, 1999, Kumar *et al.*, 2017). This vascular localization of MtNAS2 was also observed in nodules. However, the cellular localization of the protein is different to what was observed in roots; in nodules, most of vascular MtNAS2 was confined to the endodermis. This alternative distribution of MtNAS2 in vessels could be indicative of differential functions. Root vascular localization could indicate a role in metal loading of the vascular fluids, while endodermal localization in nodules might mediate either uptake from saps or intracellular metal trafficking. In any case, it seems unlikely that the nicotianamine synthesized by nodule endodermal cells would end up in the apoplast, since citrate-iron complexes seem to be formed in this compartment at a pH that does not facilitate iron-nicotianamine association (Rellán-Álvarez *et al.*, 2008).

*MtNAS2* expression in nodule core cells in the interzone and zone III also indicates a role of nicotianamine in metal homeostasis of nitrogen fixing cells. It has been previously described that nicotianamine can participate in intracellular metal trafficking and in cell-to-cell metal delivery, as well as serve as intracellular storage of metals (Haydon *et al.*, 2012). Mutation of *MtNAS2* did not significantly alter iron, copper, or zinc levels in any of the plant organs analysed, but a major shift in iron distribution was observed in nodules, with a significant decrease of iron accumulation in the interzone and early fixation zone. This would indicate that iron trafficking in these cells is altered. However, MtNAS2-mediated iron trafficking would only affect a subset of the nodule iron-proteome, since delivery to the fixation zone was not completely blocked as attested by the red colour of nodules, indicative that leghemoglobin (an important iron sink) was being produced in addition to a residual nitrogenase activity. This could suggest the existence of differential metallation pathways in nodules that might serve different subsets of proteins, which could partially complement each other under stress conditions. Supporting this hypothesis, mutation of *MtNAS2* did not equally affect all the iron species in the fixation zone. While the percentage of iron-sulfur complexes detected by XANES was significantly lower than in control plants, iron coordinated by nitrogen or oxygen atoms was increased. Considering the high demand for iron-sulfur clusters for nitrogenase assembly (Rubio and Ludden, 2005), its decrease could explain the reduction of nitrogenase activity observed. The changes in iron speciation were particularly severe in the fixation zone, which is consistent with *MtNAS2* distribution, with the observed reduction of nitrogenase activity, and with the iron distribution data. It is important to indicate that we cannot rule out similar effects on copper or zinc speciation and distribution, since the synchrotron setup available to us at the European Synchrotron Radiation Facility prevented us to carry out similar analyses on those two elements.

This work highlights the importance of MtNAS2 in iron delivery for symbiotic nitrogen fixation. This is not the only NAS gene that might be involved in the process, since total nicotianamine production is sustained in nodules, and other family members have been shown to be expressed in these organs, such as MtNAS1 (Avenhaus *et al.*, 2016). The localization of MtNAS2 indicates that nicotianamine would be involved in intracellular iron trafficking that is highly important for nitrogenase functioning. This role would not be directly providing the element to the bacteroid, since iron-citrate seems to be the key here, but perhaps would shuttle this element in the cytosol. However, to better define this possibility, new tools in elemental imaging and speciation with higher resolution within a cell need to be established to track iron and other elements. In addition, the roles of ZIF-like (Haydon *et al.*, 2012)and YSL proteins (Waters *et al.*, 2006) in symbiotic nitrogen fixation must be determined. Finally, other NAS proteins might facilitate iron recycling in *M. truncatula* nodules, as it occurs in *L. japonicus* (Hakoyama *et al.*, 2009).

## EXPERIMENTAL PROCEDURES

### Biological material and growth conditions

*M. truncatula* Gaertn R108 and *nas2-1* (NF15101) seeds were scarified in concentrated sulfuric acid (96%) for 7.5 min. After removing the acid, the seeds were washed eight times with cold water, and surface-sterilized with 50 % (v/v) bleach for 90 s. Seeds were embedded overnight in the dark at room temperature in sterile water, and transferred to 0.8 % water-agar plates for 48 h at 4 ºC (stratification). Germination was carried out at 22 ºC in the dark. Seedlings were planted on sterile perlite pots, and inoculated with *S. meliloti* 2011 or the same bacterial strain transformed with pHC60 (Cheng and Walker, 1998). Plants were grown in a greenhouse under 16 h light / 8 h dark at 25 ºC / 20 ºC conditions. In the case of perlite pots, plants were watered every two days with Jenner’s solution or water alternatively (Brito *et al.*, 1994). Nodules were obtained at 28 dpi. Plants growing in non-symbiotic conditions were watered every two weeks with a nutrient solution supplemented with 20 mM NH_4_NO_3_. For hairy-root transformation experiments, *M. truncatula* seedlings were transformed with *Agrobacterium rhizogenes* strain ARqua1, fused to the appropriate binary vector as described (Boisson-Dernier *et al.*, 2001).

### RNA Extraction and Quantitative real-time RT-qPCR

RNA was extracted from 28 dpi plants using TRI-reagent (Life Technologies), treated using DNase turbo (Life Technologies), and cleaned with RNeasy Mini-kit (Qiagen). cDNA was obtained from 500 ng RNA using PrimeScript RT reagent Kit (Takara). Expression studies were carried out by real-time reverse transcription polymerase chain reaction (RT-qPCR; StepOne plus, Applied Biosystems) using the Power SyBR Green master mix (Applied Biosystems). The primers used are indicated in Supp. Table 2. RNA levels were normalized by using the ubiquitin conjugating enzyme E2 (*Medtr7g116940*) gene as internal standard. Real time cycler conditions have been previously described (González-Guerrero *et al.*, 2010).

### GUS Staining

*MtNAS2* promoter region was obtained by amplifying the 1940 bp upstream of the start codon using the primers indicated in Supp.Table 2, and cloned by Gateway cloning technology (Invitrogen) in pDONR207 (Invitrogen) and transferred to destination vector pGWB3 (Nakagawa *et al.*, 2007). Hairy-root transformations of *M. truncatula* seedlings were carried out with *A. rhizogenes* ARqua1 as described by Boisson-Denier *et al*. (2001). After three weeks on Fahreus media plates with kanamycin (50 μg/mL), plant transformants were transferred to sterilized perlite pots and inoculated with *S. meliloti* 2011. GUS activity was determined in 28 dpi plants as described (Vernoud *et al.*, 1999).

### Confocal microscopy

The coding sequence region of *MtNAS2* and 1940 bp upstream of its start codon were cloned in pGWB13 vector (Nakagawa et al., 2007) using Gateway cloning technology (Invitrogen). This fuses three HA epitopes to C-terminus of the protein. Hairy-root *M. truncatula* transformants were transferred to sterilized perlite pots and inoculated with *S. meliloti* 2011 containing the pHC60 plasmid that constitutively expresses GFP. Nodules and roots were collected from 28 dpi plants and fixed at 4 ºC overnight in 4 % para-formaldehyde and 2.5 % sucrose in phosphate-buffered saline (PBS). Fixative was removed by washing for 5 min in PBS and 5 min in water. Nodule and roots were included in 6 % agarose for sectioning with a Vibratome 1000 Plus. Sections were dehydrated by serial incubation with methanol (30 %, 50 %, 70 % and 100 % in PBS) for 5 min and then rehydrated following the same methanol series in reverse order. Cell wall permeabilization was carried out by incubation with 2 % (w/v) cellulase in PBS for 1 h and 0.1 % (v/v) Tween 20 for 15 min. Sections were blocked with 5% (w/v) bovine serum albumin in PBS and then incubated with 1:50 anti-HA mouse monoclonal antibody (Sigma) in PBS at room temperature for 2 h. Primary antibody was washed three times with PBS for 15 min and subsequently incubated with 1:40 Alexa 594-conjugated anti-mouse rabbit monoclonal antibody (Sigma) in PBS at room temperature for 1 h. Secondary antibody was washed three times with PBS for 10 min. DNA was stained using DAPI. Images were obtained with a confocal laser-scanning microscope (Leica SP8) using excitation light at 488 nm to GFP and 561 nm for Alexa 594.

### Acetylene reduction assays

Nitrogenase activity assay was measured by acetylene reduction test (Hardy *et al.*, 1968). Wild-type and *nas2-1* nodulated roots from 28 dpi were separately introduced in 30 ml vials. Each tube contained four or five independently transformed plants. Three milliliters of air from each bottle was replaced by the same volume of acetylene, tubes were subsequently incubated for 30 min at room temperature. Gas samples were measured by analyzing 0.5 ml of ethylene from each bottle in a Shimadzu GC-8A gas chromatograph using a Porapak N column. The amount of the ethylene produced was determined by measuring the ethylene peaks relative to the standards.

### Chlorophyll Content Assays

Total chlorophyll content was determined as previously described with some modifications (Inskeep and Bloom, 1985). Leaves were collected from 28 dpi plants and poled to obtain 50 mg of fresh material. Chlorophyll was extracted with 500 μl of di-methyl-formamide at 4 ºC overnight. Leaves were centrifuged for 5 min at 600 g at room temperature. After transferring the supernatant to another vial, the chlorophyll extraction was repeated with the same leave using strong vortexing. After spinning for 5 min at 600 g, the supernatant was pooled with the previous one. Chlorophyll was quantified at 647 nm and 664 nm in a Ultraspec 3300 spectrophotometer (Amershan Bioscience).

### Metal content determination

Shoots, roots, and nodules were collected from 28 dpi plants and mineralized with 15.6 M HNO_3_ (trace metal grade) at 75°C for 3 h and 2 M H_2_O_2_ at 20°C overnight. Metal quantifications were performed in duplicate by Atomic Absorption Spectroscopy, using a Perkin Elmer PinAAcle 900Z GF-AAS equipment. Metal concentration was normalized against fresh tissue weight.

### Synchrotron Radiation X-Ray Fluorescence Spectroscopy (XRF) and XANES

XRF hyperspectral images and μXANES spectra were acquired on the beamline ID21 of the European Synchrotron Radiation Facility (Cotte *et al.*, 2017), at 110 K in the liquid nitrogen (LN2) cooled cryostat of the Scanning X-ray Micro-spectroscopy end-station. Seven sections from *M. truncatula* R108 nodules and five from *nas2-1* nodules were obtained from different nodules embedded in OCT medium and cryo-fixed by plunging in isopentane chilled with LN2. The 25 μm-thick sections of frozen samples were obtained using a Leica LN2 cryo-microtome and accommodated in a Cu sample holder cooled with LN2, sandwiched between Ultralene (SPEX SamplePrep) foils. The beam was focused to 0.4×0.9 μm^2^ with a Kirkpatrick-Baez (KB) mirror system. The emitted fluorescence signal was detected with energy-dispersive, large area (80 mm^2^) SDD detector equipped with Be window (SGX from RaySpec). Images were acquired at the fixed energy of 7.2 keV, by raster-scanning the sample in the X-ray focal plane, with a step of 3×3 μm^2^ and 100 ms dwell time. Elemental mass fractions were calculated from fundamental parameters with the PyMCA software package, applying pixel-by-pixel spectral deconvolution to hyperspectral maps normalized by the incoming flux (Solé *et al.*, 2007). The incoming flux was monitored using a drilled photodiode previously calibrated by varying the photon flux at 7.2 keV obtaining a response of 1927.9 charges/photon with a linear response up to 200 kcps. In PyMCA the incoming flux and XRF detector parameters were set to 2×10^9^ photons/s, 0.7746 cm^2^ active area, and 4.65 cm sample to XRF detector distance. Sample matrix was assumed to be amorphous ice (11% H, 89% O, density 0.92 g/cm^3^), the sample thickness set at 25μm obtained with the use of a cryo-microtome.

Fe-K edge (7.050 to 7.165 keV energy range, 0.5 eV step) μXANES spectra were recorded in regions of interest of the fluorescence maps acquired on ID21 beamline. Individual spectra were processed using Orange software with the Spectroscopy add-on (Demsar *et al.*, 2013). The pre-processing step consisted of vector normalization and a Savitsky-Golay filter for the smoothing. Then a principal component analysis was performed on the second derivative of the spectra to highlight potential differences among genotypes within a given region of the nodule. A reference library was used for linear combination fitting (LCF) procedure. This library consisted of: Fe-foil (Fe(0)), Fe(II)-nicotianamine, Fe(II)S2 (http://ixs.iit.edu/database/), Fe(III)-haem (50 mM, pH7, bought from Sigma, CAS number: 16009-13-5), Fe(III)-cellulose (5 mM FeCl_3_ + 50 mM cellulose, pH 5.8, bought from Sigma, CAS number: 9004-34-6), Fe(III) glutamic acid (5 mM FeCl_3_ + 50 mM glutamic acid, pH 7, bought from Sigma, CAS number: 56-86-0) and Fe(III) ferritin (bought from Sigma, CAS number: 9007-73-2, 50 mM, pH7).Reference compounds were classified as Fe(II)-S (FeS2), Fe(II)-O/N (Fe-NA) and Fe(III)-O (Fe-cellulose, Fe-glutamic acid and ferritin). XANES data treatment was performed using Athena software (Ravel and Newville, 2005) as previously described (Larue *et al.*, 2014).

### Statistical tests

Data were analyzed by Student’s unpaired *t* test to calculate statistical significance of observed differences. Test results with *p*-values < 0.05 were considered as statistically significant.

## Acknowledgments

This research was funded by a European Research Council Starting Grant (ERC-2013-StG-335284) and a Ministerio de Economía y Competitividad (MINECO) grant (AGL2015-65866-P), to MGG, and a MINECO grant (AGL2016-75226-R) to JA and AA-F. VE was partially funded by the Severo Ochoa Programme for Centres of Excellence in R&D from Agencia Estatal de Investigación of Spain (grant SEV-2016-0672) to CBGP. IA is recipient of a Juan de la Cierva-Formación postodoctoral fellowship from Ministerio de Ciencia, Innovación y Universidades (FJCI-2017-33222). Development of *M. truncatula Tnt1* mutant population was, in part, funded by the National Science Foundation, USA (DBI-0703285) to KSM. We would like to thank Dr. Marine Cotte and Dr Juan Herrera-Estrella for assistance in using beamline ID21 during experiments EV246 and EV323. We would also like to acknowledge the other members of laboratory 281 at Centro de Biotecnología y Genómica de Plantas (UPM-INIA) for their support and feedback in preparing this manuscript.

## SHORT LEGEND FOR SUPPORTING INFORMATION

**Supporting Figure 1.** Anatomy of 28 dpi wild-type and *nas2-1* nodules.

**Supporting Figure 2.** Nicotianamine content in wild-type and *nas2-1* plants.

**Supporting Figure 3.** Phenotype of *nas2-1* plants under low iron conditions.

**Supporting Figure 4.** Control for immunolocalization assays.

**Supporting Table 1.** Genes affected by *Tnt1* insertion in *M. truncatula* line NF15101.

**Supporting Table 2.** Primers used in this study.

**Supporting Materials and Methods**

