## Supplementary material for "Nicotianamine synthase 2 is required for symbiotic nitrogen fixation in *Medicago truncatula* nodules": Supp.

**Supporting Table 1.** *Tnt1* gene insertions in *M. truncatula* NF15101 mutant line. N.R stands for non-reported.

| Insertion | Gene ID | Homology to |
|---|---|---|
| high 1 | N.R. | |
| high 2 | N.R. | |
| high 3 | *Medtr2g079280* | ubiquitin-conjugating enzyme |
| high 4 | N.R. | |
| high 5 | N.R. | |
| high 6 | *Medtr5g023730* | fatty acyl-CoA synthetase family protein |
| high 7 | *Medtr0400s0040* | LRR receptor-like kinase family protein |
| high 8 | NOL1/NOP2/sun family protein | Coexpressed with genes in leaf specific coexpression subnetwork |
| high 9 | N.R. | |
| high 10 | N.R. | |
| high 11 | *Medtr5g064610* | Thylakoid lumenal 29 kDa protein |
| high 12 | N.R. | |
| high 13 | *Medtr8g038400* | ADP-ribosylation factor GTPase-activating protein AGD10 |
| high 14 | *Medtr5g016060* | phototropic-responsive NPH3 family protein |
| high 15 | *Medtr5g021860* | F-box protein interaction domain protein |
| high 16 | N.R. | |
| high 17 | N.R. | |
| high 18 | N.R. | |
| high 19 | *Medtr2g095800* | xyloglucan endotransglucosylase/hydrolase family protein |
| high 20 | *Medtr2g070310* | MtNAS2 |
| high 21 | N.R. | |
| high 22 | N.R. | |

Supporting Table 2. Primers used in this study

| Name | Sequence | Use |
|---|---|---|
| Fw MtNAS2P- GW | GGGGACAAGTTTGTACAAAAAAGCAGGCTCTAA AATTGACAATTGTTATCA | Cloning of *MtNAS2* promoter in Gateway vector |
| Rv MtNAS2P-GW | GGGGACCACTTTGTACAAGAAAGCTGGGTATGG ATTGAATTGAAGTAATTT | Cloning of *MtNAS2* promoter in Gateway vector |
| Rv MtNAS2 GW | GGGGACCACTTTGTACAAGAAAGCTGGGTAAGCT TTAGGTTTGCGAAGAAG | Cloning of *MtNAS2* gene in Gateway vector |
| Fw MtNAS2 | TGAAATTCCACAAGAGCTTCTCA | qRT-PCR of *MtNAS2* |
| Rv MtNAS2 | TGGATGAAGGGAGGATGCAAA | qRT-PCR of *MtNAs2* |
| Fw MtUBQ9 | GGTTGATTGCTCTTCTCTCCCC | qRT-PCR of gene encoding for ubiquitin conjugating enzyme E2 |
| Rv MtUBQ9 | AAGTGATTGCTCGTCCAACCC | qRT-PCR of gene encoding for ubiquitin conjugating enzyme E2 |

### Supplemental Materials and Methods

Nodule structure analyses

Sections for nodule structural organization were obtained from 28 days-post-inoculation (dpi) nodules fixed in 1 % formaldehyde, 0.5 % glutaraldehyde, 2.5% sucrose in 50 mM potassium phosphate buffer (pH 7.4) at 4 °C for 14 – 16 h in gentle agitation. Samples were dehydrated with ethanol series and embedded in LR-White resin (London Resin Company Ltd, UK). Nodule were placed in gelatine capsules, filled with LR-White resin and polymerized at 60 °C for 24 h. Serial thin section of 1 µm, were cut using a diamond knife in a Reichert Ultracut S-Ultramicrotrome (Leyca), stained with a mixture of 1 % toluidine blue in aqueous 1 % di-sodium tetra-borate in water Direct observation of sections was perfected by a Zeiss Axiophot photomicroscope (Carl Zeiss, Oberkochen, Germany) with pictures taken with a digital camera (Leica DFC 420C, Leica).

Nicotianamine concentration

Nicotianamine was extracted as previously described (Banakar et al., 2017) with some modifications. Briefly, nicotianamine was extracted from approximately 200 mg of roots and shoots or from 50 mg of nodules, and frozen grinded in 400 µl miliQ water spiked with nicotyl-lysine (9 µM final concentration) as internal standard. Samples were homogenized in a mixer mill (Retsch MM300, Retsch) during 5 minutes at $30^{-1}$ sec frequency, and then centrifuged at 15000 g for 30 minutes at 4 ºC. Supernatant was then passed through a 3 kDa cutoff centrifugal filter (cellulose Amicon® , Merck) and dried under vacuum. Dry residues from shoots were dissolved in 20 µl of miliQ water, whereas root-and-nodule dry residues were dissolved in 12 µl. Then 5 µl aliquots were mixed with EDTA (final concentration 8.33 mM) to dissociate potential nicotianamine-metal complexes, and 50% (v/v) mobile phase A (see below) to favor chromatographic separation. The mixture was filtered through 0.45 polyvinylidene fluoride (PVDF) ultrafree-MC centrifugal filter devices (Merck) before analysis. Nicotianamine (Toronto Research Chemicals) calibration curve were prepared with nycotyl-lysine as internal standard.

Nicotianamine levels were determined by high-performance liquid chromatography electrospray ionization time-of-flight mass spectrometry (HPLC-ESI-TOF-MS) as described by Banakar et al. (2017). The samples were fractionated using an Alliance 2795 HPLC system (Waters) and µLC column (SeQuant ZIC®-HILIC, 15 cm x 1 mm internal diameter, 5 µm, 200 Å, Merck), with a mobile phase consisting of solvent A (9:1 10 mM ammonium acetate:acetonitrile, pH 7.3) and solvent B (8:2 30 mM ammonium acetate:acetonitrile, pH 7.3) at a flow rate of 0.15 mL $min^{-1}$. The gradient program started at 100% (v/v) solvent A for 3 min, and then decreased linearly to 30% (v/v) solvent A over the next 7 min, then remained for 7 min at 30% (v/v) solvent A, and then returned to the initial conditions over the next 8 min. The column was then allowed to stabilize for 10 min at the initial conditions before proceeding to the next injection. The total HPLC run time was 35 min, the injection volume was 10 µl and the auto sampler and column temperatures were 6 ºC and 30 ºC, respectively. The HPLC was coupled to the MicrOTOF mass spectrometer (Bruker Daltonics) equipped with an ESI source. The operating conditions were optimized by the direct injection of 100 µM solutions of nicotianamine standard at a flow rate of 180 µl $h^{-1}$. Mass spectra were acquired in negative ion mode over the 150–700 mass-to-charge (*m/z*) ratio range. The mass axis was calibrated externally using Li–formate adducts (10 mM LiOH, 0.2% (v/v) formic acid and 50% (v/v) 2-propanol). Bruker Daltonik software packages micrOTOF Control v2.2, HyStar v3.2 and Data Analysis v4.0 were used to control the MS, HPLC interface and for data processing, respectively. Nicotianamine concentrations were quantified by external calibration, using nicotyl-lysine as internal standard.

**SUPPL. REFERENCE**

**Banakar R, Álvarez-Fernández A, Abadía J, Capell T, Christou P.** (2017) The expression of heterologous Fe (III) phytosiderophore transporter *HvYS1* in rice increases Fe uptake, translocation and seed loading and excludes heavy metals by selective Fe transport. *Plant Biotechnol. J.* **15**, 423-432.

SUPPORTING FIGURE LEGENDS

**Supporting Figure 1.** Anatomy of 28 days-post-inoculation (dpi) wild type (left panel) and *nas2-1* (right panel) nodules. Toluidine blue stain of sections of 28 WT and *nas2-1* nodules. Bars = 50 μm.

**Supporting Figure 2.** Nicotianamine concentration in wild type (WT) and *nas2-1* 28 dpi plants. Data are the mean ± SE of n = 5-8 samples from two independent experiments.

**Supporting Figure 3.** Phenotype of *nas2-1* plants under low iron conditions. (A) Growth of representative wild type (WT) and *nas2-1* plants when watered with a nutrient solution supplemented with ammonium nitrate and not inoculated with *S. meliloti*. Bar = 1.5 cm. (B) Fresh weight of WT and *nas2-1* plants. Data are the mean ± SE of at least 5 plants. (C) Chlorophyll concentration of wild type and *nas2-1* plants. Data are the mean ± SE of two sets of 5 pooled plants.

**Supporting Figure 4.** Control for immunolocalization assays. (A) Longitudinal section of a 28 days-post-inoculation (dpi) *M. truncatula* nodule expressing *MtNAS2-HA* under its own promoter. Sections were treated as indicated for confocal microscopy, but without adding the Alexa594 conjugated antibody. The left panel corresponds to the Alexa594 emission signal. Transformed plants were inoculated with a GFP-expressing *S. meliloti* (green, middle panel). Both channels were overlaid with the transillumination image (right panel). Bars = 100 μm. (B). Cross section from a 28 dpi *M. truncatula* root expressing *MtNAS2-HA* under its own promoter. Left panel corresponds to the Alexa594 signal used to detect the HA-tag, middle panel corresponds to autofluorescence signal of xylem, and the two were overlaid with the bright field channel in the right panel. Bars = 50 μm.

**SUPPORTING FIGURE 1**

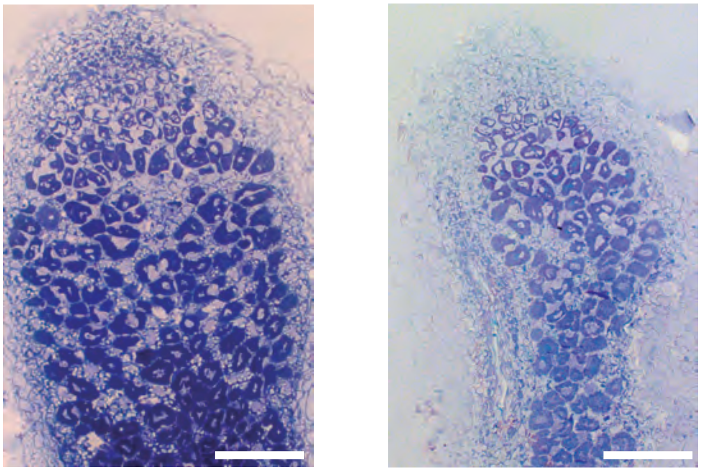

**SUPPORTING FIGURE 2**

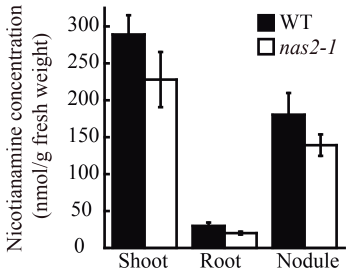

### SUPPORTING FIGURE 3

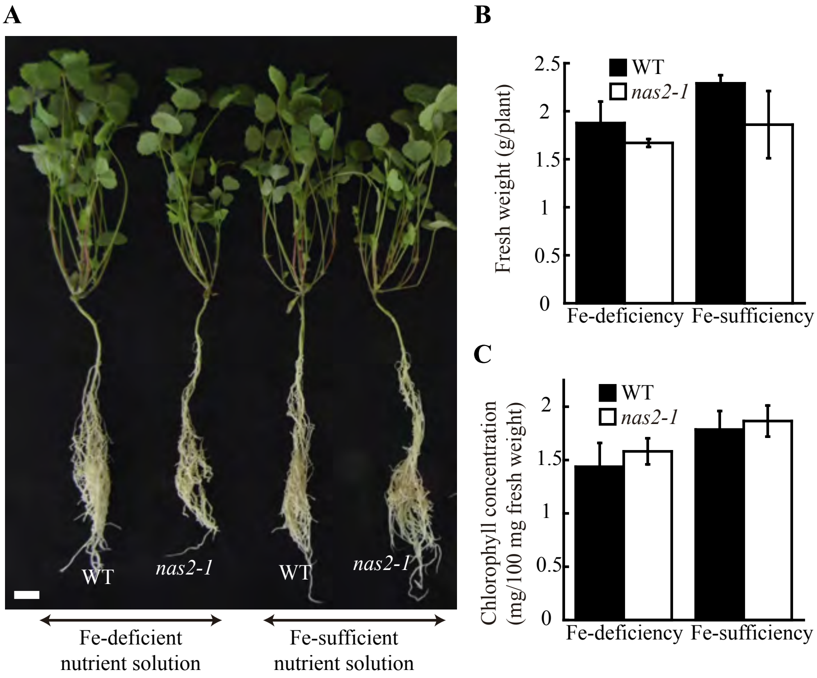

### SUPPORTING FIGURE 4

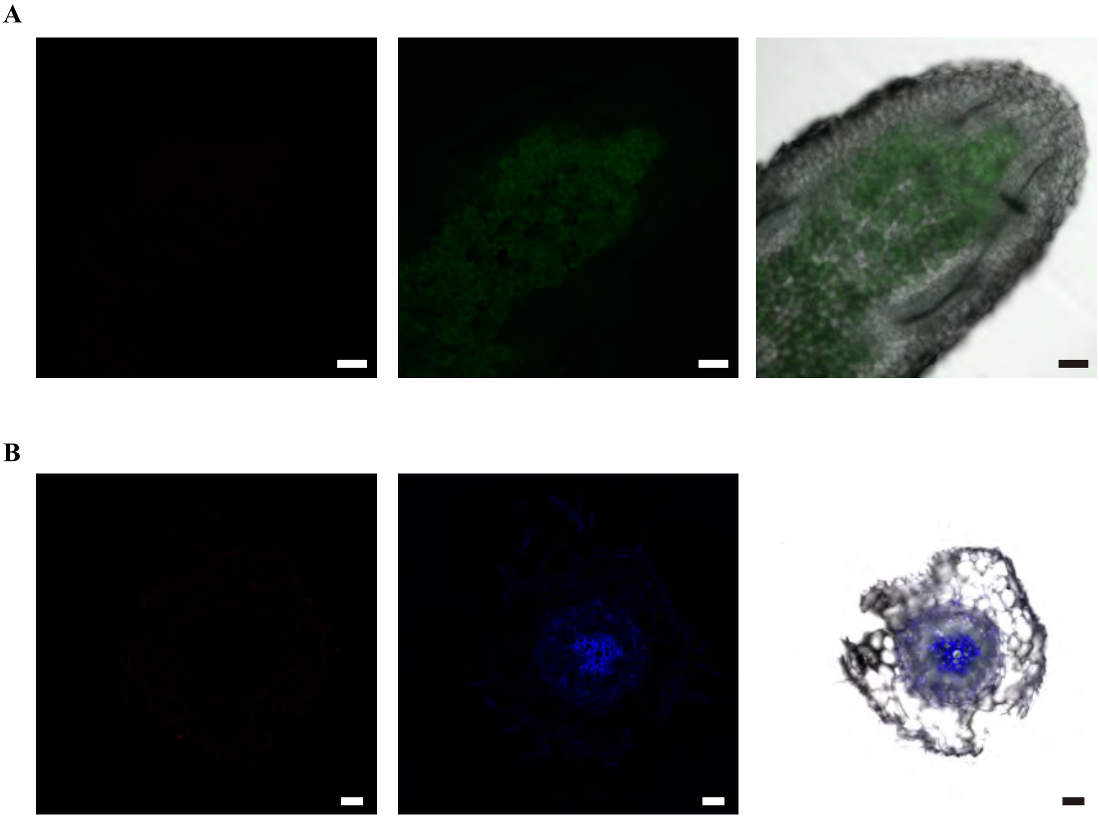
